# Differential transcriptional profiles of vagal sensory neurons in female and male mice

**DOI:** 10.1101/2024.02.15.580501

**Authors:** Young-Hwan Jo

## Abstract

The differences in metabolic homeostasis, diabetes, and obesity between males and females are evident in rodents and humans. Vagal sensory neurons in the vagus nerve ganglia innervate a variety of visceral organs and use specialized nerve endings to sense interoceptive signals. This visceral organ-brain axis plays a role in relaying interoceptive signals to higher brain centers as well as in regulating the vago-vagal reflex. I hypothesized that molecularly distinct populations of vagal sensory neurons would result in differences in metabolic homeostasis between the sexes. Single-nucleus RNA sequencing analysis of vagal sensory neurons from females and males reveals differences in the transcriptional profiles of cells in the vagus nerve ganglia. These differences are linked to the expression of sex-specific genes such as *Xist, Tsix*, and *Ddx3y*. Among the 13 neuronal clusters, one-fourth of the neurons in male mice are located in the *Ddx3y*-enriched VN1 and VN8 clusters, which display a higher enrichment of *Trpv1, Piezo2, Htr3a*, and *Vip* genes. In contrast, 70% of the neurons in females are found in *Xist*-enriched clusters VN4, 6, 7, 10, 11, and 13, which show enriched genes such as *Fgfr1, Lpar1, Cpe, Esr1, Nrg1, Egfr*, and *Oprm1*. Two clusters of satellite cells are identified, one of which in males contains some oligodendrocyte precursor cells. A small population of cells express *Ucp1* and *Plin1*, indicating that they are epineural adipocytes. Understanding the physiological consequences of differences in these transcriptomic profiles on energy balance and metabolic homeostasis would help develop sex-specific treatments for obesity and metabolic dysregulation.

## Introduction

The visceral organ-brain axis is essential for maintaining metabolic homeostasis and other vital functions such as cardiovascular function, breathing, digestion, food intake, and reward ^1-7^. Vagal sensory neurons located in the vagus nerve ganglia consisting of the jugular and nodose ganglia transmit interoceptive signals from visceral organs to the medulla via the vagus nerve. Recent advances in neural mapping and single-cell RNA sequencing have enabled to identify vagal sensory neurons innervating diverse visceral organs, including the larynx, stomach, intestine, pancreas, heart, and lung ^1-4,7-10^. These prior studies have revealed the highly specialized cellular, molecular, and anatomical organization of vagal sensory neurons ^1-4,7-12^. For instance, single-cell RNA-seq analysis has uncovered 12-37 molecularly unique subtypes of vagal sensory neurons ^1,6,11^. Furthermore, molecularly distinct subsets of vagal sensory neurons have specific nerve terminals depending on their sensory modalities and functions ^1,2,4,10^, and their functional properties are largely determined by their molecular identity ^1-4,9,10^.

It is widely recognized that there are significant differences in metabolic homeostasis, diabetes, and obesity between males and females, both in rodents and humans. Female rodents are less susceptible to diet-induced obesity, insulin resistance, hyperglycemia, and hypertriglyceridemia ^13-19^. Similarly, premenopausal women are generally protected from metabolic diseases ^20^, whereas men and postmenopausal women are more likely to accumulate fat in the intra-abdominal depot, which increases the risk of developing obesity-related metabolic complications ^19^. Sex hormones, particularly estrogen, play a significant role in causing this difference in metabolic physiology between the sexes ^15,20,21^. In addition to the contribution of sex hormones to metabolic homeostasis, it is plausible that molecularly distinct types of vagal sensory neurons may differentially integrate interoceptive signals at the peripheral level and transmit this filtered information to higher brain centers in a sex-dependent manner. In other words, the expression of genes encoding membrane receptors, ion channels, neurotransmitters, neuropeptides, and intracellular signaling proteins in vagal sensory neurons would be sex- and cell-specific. In this study, I sought to identify differences in the transcriptional profiles of cells in the vagus nerve ganglia of male and female mice.

## Results

### Heterogeneity of vagal sensory neurons in mice

SnRNA-Seq was conducted on dissociated cells from the vagus nerve ganglia using the 10X Genomics Chromium platform (Figure 1A). Low-quality cells and possible doublets were removed using the Cell Ranger and DIEM programs. I analyzed a total of 8,714 cells (3791 cells from females and 4923 cells from males). The t-distributed stochastic neighbor embedding (t-SNE) projection revealed 20 individual clusters (Figure 1B and C and Supplementary. Table 1). Of the 20 clusters, 13 clusters expressed neuronal marker genes, including microtubule associated protein 2 (*Map2)*, synapsin 1 and 2 (*Syn1 and Syn2*), and solute carrier family 17 member 6 (*Slc17a6*), indicating that they were vagal sensory neurons (VN; n= 6,160 out of 8,714 cells). These neurons were further subdivided into 13 clusters (VN1-VN13). Among the 20 clusters, three expressed glial cell marker genes: periaxin (*Prx*), non-compact myelin-associated protein (*Ncmap*), and myelin protein zero (*Mpz*) (Figure 1D). One of these clusters was identified as myelinating Schwann cells based on the high expression of peripheral myelin protein 2 (*Pmp2*), ADAMTS-like 1 (*Adamtsl1*), and Erb-B2 receptor tyrosine kinase 4 (*Erbb4*) compared to the other two clusters (Supplementary Table 2). These genes are considered myelinating Schwann cell markers ^22,23^. Only one cluster expressed mannose receptor c-type 1 (*Mrc1*), a microglia (MG) marker gene (n=267 cells). Additionally, two clusters expressed endothelial cell (EC) marker genes: platelet-derived growth factor receptor alpha (*Pdgfra*) and endomucin (*Emcn*) (Figure 1D). Interestingly, a small number of cells expressed adipocyte marker genes, including perilipin 1 (*Plin1*), uncoupling protein 1 (*Ucp1*), and peroxisome proliferator-activated receptor gamma (*Pparg*) (Figure 1C and D), suggesting that they were epineural adipocytes (EA; n=132 cells).

**Figure 1.**
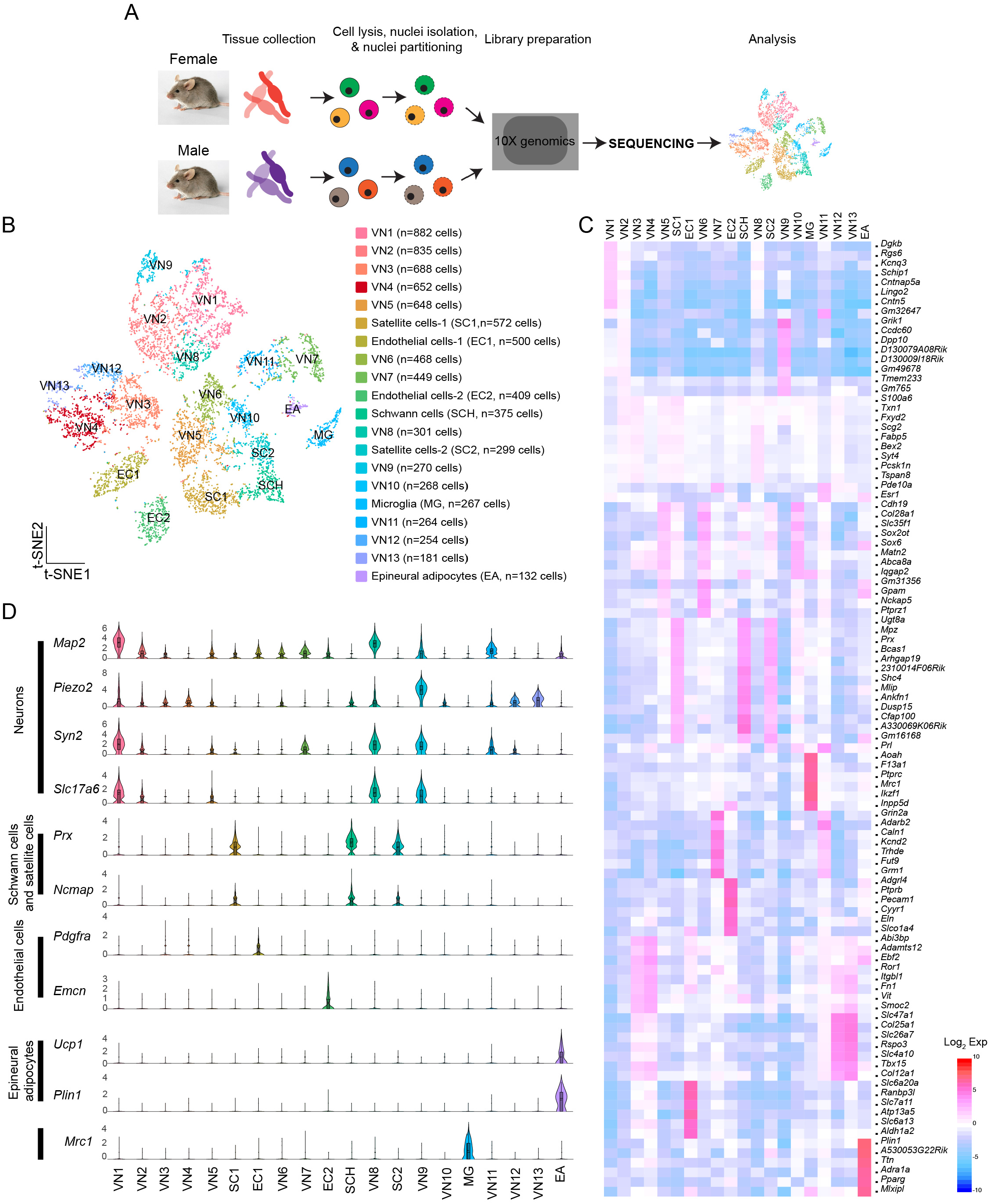
Molecular heterogeneity of cells in the vagus nerve ganglia. **A**. Schematic illustration of snRNA-Seq workflow **B**. t-SNE projection showing twenty unique clusters, comprised of thirteen neuronal clusters, three glial cell clusters, one microglia cluster, and one epineural adipocytes group. **C**. Heatmap plot showing differential gene expression among twenty clusters. **D**. Violin plots displaying the enriched genes specific to each cell type within each specific group.

### Sex differences in the transcriptional profiles of vagal sensory neurons

I then sought to determine if there are sex differences in the transcriptional profiles of the cells in the vagus nerve ganglia. As expected, X-linked X-inactive-specific transcript (*Xist*) RNA, which plays a crucial role in regulating X-chromosome inactivation ^24^, was present in female mice (Fig. 2A). On the other hand, the DEAD-box helicase 3 Y-linked (*Ddx3y*) gene, which is located on the Y chromosome, was detected in male mice (Fig. 2A). Unsupervised uniform manifold approximation and projection (UMAP) plots revealed that there were 20 clusters for both males and females, but there were noticeable differences in the number of cells in the same clusters between females and males (Fig 2B and C). In other words, the number of cells in the same clusters was not evenly distributed, with females having more cells in VN4, VN7, VN10, and VN13, whereas males had more cells in VN1, VN2, VN3, VN5, VN8, and VN12 (Fig. 2D). Additionally, males had a higher number of cells in SC1, whereas females had a greater number of cells in SC2. There was a significant difference in the number of satellite cells (Fisher’s exact test, p<0.001) and Schwann cells (Fisher’s exact test, p<0.001) between the sexes.

**Figure 2.**
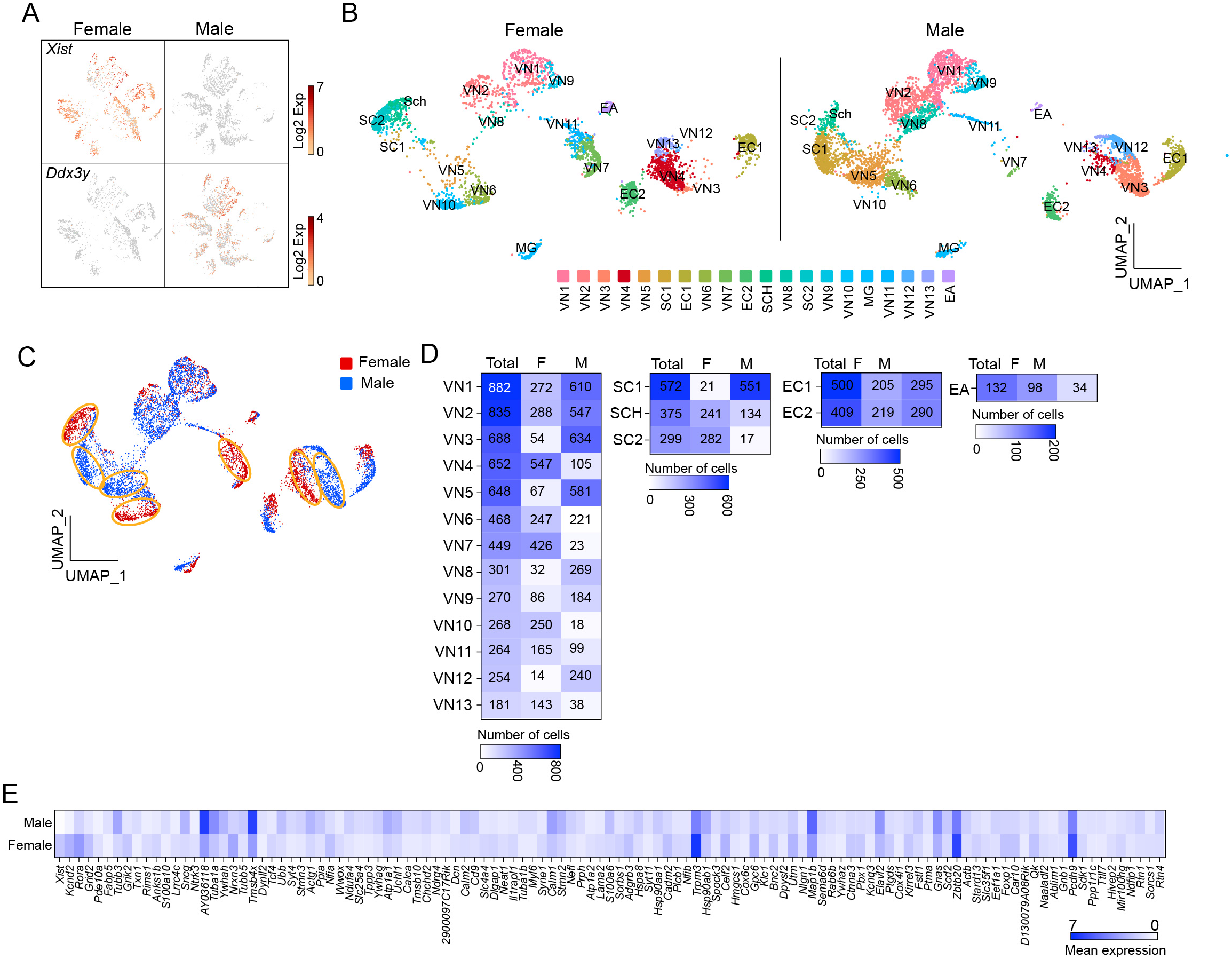
Uneven distribution of the number of cells within the same clusters. **A**. t-SNE projections highlighting the expression of *Xist* and *Ddx3y* in females and males, respectively. **B** and **C**. UMAP projections showing twenty molecularly distinct clusters in females and males. Panel C illustrates a clear difference in the number of cells in the same clusters between females and males. **D**. Heatmap plots showing the number of cells in each cluster between females and males. **E**. Heatmap plot displaying the differential expression of genes between males and females

Differential gene expression analysis revealed that more than 100 genes differed significantly between females and males (Fig. 2E and Supplementary. Table 3). These included genes encoding voltage-dependent channels, ligand-gated channels, membrane receptors, intracellular signaling proteins, and transcription factors (i.e., *Kcnd2, Kcnq3, Grid2, Grik2, Rora, Gnas, Pde10a, Ndrg4, Nfia, and Tcf4*)(Supplementary. Table 3). Upon conducting a selective gene ontology (GO) analysis using GOrilla ^25^ of the significantly upregulated genes in male and female mice, it was observed that among the top 20 enriched GO terms in biological process, they shared eight GO terms related to cell development and differentiation (Supplementary. Fig. 1). Interestingly, one-fourth of the enriched GO terms in females were related to synaptic signaling, whereas one-third of them in males were associated with intracellular component organization and transport (Supplementary. Fig. 1).

I further evaluated whether the uneven distribution of cell numbers in the same clusters is associated with sex differences. Gene distribution plots revealed that both *Xist* and X (inactive)-specific transcript (*Tsix*) that binds *Xist* during X chromosome inactivation ^26^ were expressed in VN4, VN6, VN7, VN10, VN11, and VN13 (Fig. 3A). In contrast, *Ddx3y* was enriched only in VN1 and VN8 (Fig. 3A). In males, the percentage of cells in VN1 and VN8 was 25% (n= 879 out of 3,569 neurons), whereas in females, it was 12% of the total number of neurons (n=301 out of 2,591 neurons). In contrast, females had a significantly higher proportion of VN4, VN6, VN7, VN10, VN11, and VN13, accounting for 69% of the cells in the neuronal clusters, compared to only 18% in males. VN9 expressed both *Xist* and *Ddx3y*, while the expression of the three genes was very low or minimal in VN2, VN3, VN5, and VN12 (Fig. 3A). The UMAP plots further demonstrated the uneven distribution of the number of cells in the same clusters (Fig. 3B).

**Figure 3.**
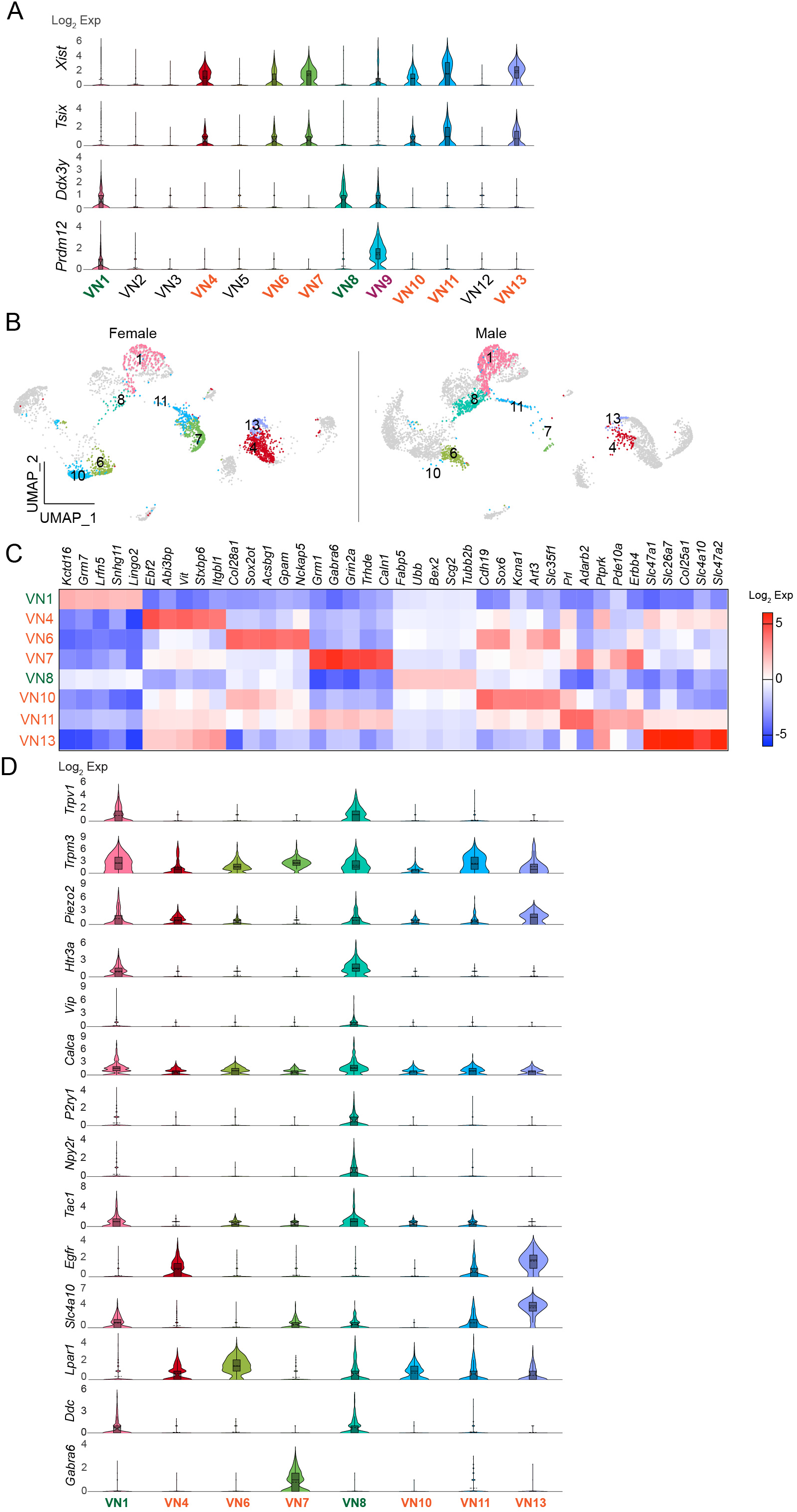
Existence of sex-specific neuronal clusters. **A**. Violin plots depicting the differential expression of *Xist, Tsix*, and *Ddx3y* in each neuronal cluster. VN9 showed expression of *Prdm12*. **B**. UMAP projections showing *Xist*- and *Ddx3y*-enriched neuronal clusters, specifically *Xist*-enriched clusters like VN4, VN6, VN7, VN10, VN11, and VN13, and *Ddx3y*-enriched clusters like VN1 and VN7. **C**. Heatmap plot showing the top 5 genes expressed in each cluster. **D**. Violin plots displaying the enriched genes specific to *Xist*- and *Ddx3y*-positive neuronal clusters. VN1 and VN8 exhibited elevated expression such as *Trpv1, Trpm3, Piezo2, Htr3a, Calca, Tac1*, and *Ddc* compared to other neuronal clusters.

Comparison of the transcriptional profiles in the neural clusters revealed that more than one thousand genes were differentially expressed among these clusters (Supplementary Table 4). Fig. 3C shows the top five enriched genes in VN1, VN4, VN6, VN7, VN8, VN10, VN11, and VN13.

I also examined the expression of genes that are commonly present in vagal sensory neurons, specifically chemosensitive transient receptor potential cation channel subfamily V member 1 (*Trpv1*), temperature-sensitive TRP subfamily M member 3 (*Trpm3*), and mechanosensitive piezo-type ion channel component 2 (*Piezo2*) (Fig. 3D). Intriguingly, *Trpv1* was mainly found in VN1 and VN8 (Fig. 3D). Although the degree of *Trpm3* and *Piezo2* expression was different, both genes were expressed across the clusters (Fig. 3D). Molecularly distinct subsets of vagal sensory neurons have their own target organs. For example, vagal sensory neurons that control the digestive system include vasoactive intestinal peptide (VIP)-, glucagon-like peptide 1 receptor (GLP1R), oxytocin receptor (OXTR), G protein-coupled receptor 65 (GPR65), somatostatin (SST), 5-hydroxytryptamine receptor 3A (HTR3A)-, and calcitonin gene-related peptide alpha (CALCA)-expressing neurons ^1,4,10^. The purinergic receptor P2Y1 (P2RY1)- and neuropeptide Y receptor Y2 (NPY2R)-expressing neurons innervate the lungs ^9^. The pancreatic islets are innervated by vagal sensory neurons expressing tachykinin 1 (TAC1), CALCA, and HTR3 ^3^. I thus examined if there are differences in their gene distribution across the *Xist*- and *Ddx3y*-positive VN clusters. Intriguingly, VN1 and VN8 expressed genes, including *Htr3a, Vip, Calca, P2ry1, Npy2r*, and *Tac1*, although the extent of gene distribution was different (Fig. 3D). On the other hand, VN4, VN6, VN7, VN10, VN11, and VN13 displayed low or minimal expression levels of these genes (Fig. 3D). As the expression levels of *Glp1R, Oxtr, Gpr65*, and *Sst genes* were minimal, the difference could not be observed.

### Sex differences in the transcriptional profiles of satellite cells

As the number of satellite cells in SC1 and SC2 was significantly different (Fisher’s exact test, p<0.001; Fig. 2D and Fig. 4A), I examined whether this was also related to sex differences. The results showed that *Xist* expression levels were higher in SC2 than in SC1 (Fig. 4B). Moreover, There were differential gene expression between the two clusters (Fig. 4C). Tubulin beta 3 (*Tubb3*) expression was significantly higher in SC1 than in SC2. *Tubb3*, a gene typically associated with neurons, has also been discovered in intermediate cells that lie between oligodendrocyte precursor cells (OPCs) and astrocytes (28). Furthermore, several other genes, including laminin subunit alpha (*Lmna*), CNPase (*Cnp*), and SRY-box transcription factor 10 (*Sox10*), which are enriched in OPC ^27,28^, were found in SC1 (Fig. 4B). Specifically, as LMNA expression levels rise as OPCs differentiate ^28^, SC1 may consist of some progenitor cells that differentiate into myelinating Schwann cells. The number of Schwann cells was significantly lower in male mice than in female mice (Fisher’s exact test, p<0.001; Fig. 2D). In contrast, EC1 and EC2 had comparable numbers of cells (Fisher’s exact test, p>0.05; Figure 4D), and both clusters expressed *Xist* (Figure 4E).

**Figure 4.**
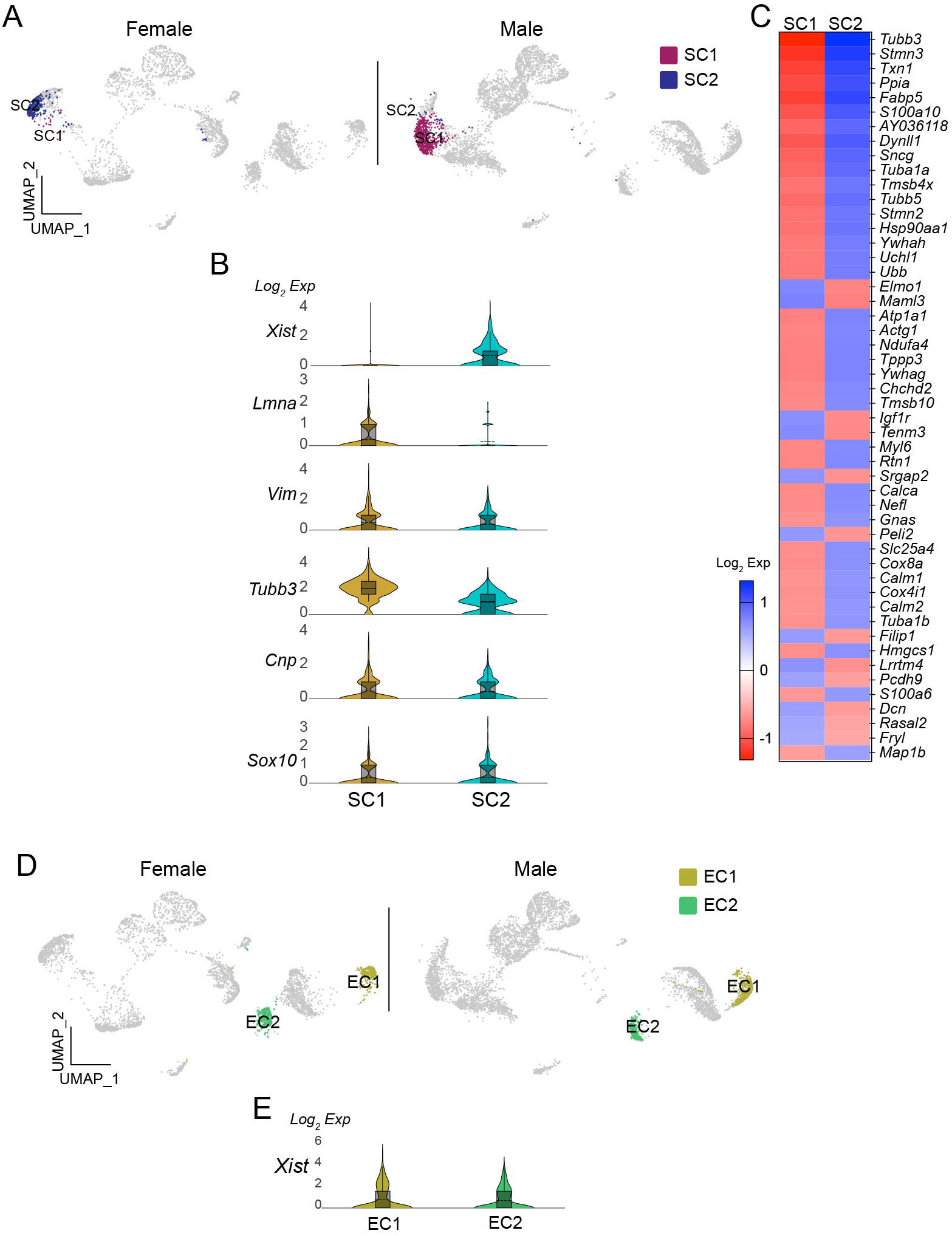
Existence of sex-specific glial cell populations in the vagus nerve ganglia. **A**. UMAP projections showing SC1 and SC2 clusters in females and males. **B**. Violin plots displaying differential expression of *Xist* and OPC-related genes in SC1 and SC2 clusters. **C**. Heatmap plot showing the differential gene expression in SC1 and SC2 clusters. **D**. There were no differences in the number of EC and *Xist* expression in females and males.

## Discussion

This study demonstrated distinct transcriptomic profiles of cells, including sensory neurons and glial cells in the vagus nerve ganglia, between male and female mice. Prior research has established the molecular and neuroanatomical identity of target organ-specific vagal sensory neurons in several studies ^1,3,4,9,10^. Although some of these studies utilized both male and female mice ^1,9,10^, they reported no sex differences without specifying which results were obtained from male and female mice. Vagal sensory neurons with sophisticated nerve terminals dependent on target areas in organs are vital for detecting and relaying interoceptive signals from organs to the brain ^1,10^. This implies that they may possess the capacity to regulate the energy balance. In fact, activating gastrointestinal mechanoreceptors has been shown to reduce food intake ^1^. Metabolic homeostasis, diabetes, and obesity exhibit notable differences between males and females in both rodents and humans ^13-20^. In this regard, differences in the quantity and gene expression profiles of cells in the vagus nerve ganglia between the sexes may respond distinctly to nutrients and environmental cues, ultimately leading to distinct metabolic regulation.

Thirteen neuronal clusters with unique molecular characteristics were identified. Approximately half of these neuronal clusters exhibited high expression levels of both *Xist* and *Tsix* genes, whereas only two showed elevated levels of the *Ddx3y* gene. One of these clusters (VN9) expressed both *Xist* and *Ddx3y* and showed high levels of *Prdm12*, a marker gene for neurons in the jugular ganglia ^11^. Another cluster (VN1) also expressed *Prdm12* and had both the jugular marker *Tac1* and the nodose marker *Htr3a* ^29^, suggesting that a small subset of VN1 neurons were jugular neurons. A comparison of genes that have been extensively studied in vagal sensory neurons revealed that VN1 and VN8 displayed a higher enrichment of *Trpv1, Piezo2, Htr3a, Vip, Calca, P2ry1, Npy2r*, and *Tac1* genes ^1,3,4,9,10^ compared to the clusters with high expression of the *Xist* and *Tsix* genes. As they expressed chemosensitive, temperature-sensitive and mechanosensitive nociceptors, they are polymodal sensory neurons and seem to be involved in various physiological processes. PIEZO2-positive neurons with mechanosensory endings played a role in the baroreceptor reflex ^30^, while activation of P2RY1- and NPY2R-positive neurons produced opposing effects on breathing; stimulation of P2RY1-positive cells silenced respiration, while activation of NPY2R neurons induced rapid, shallow breathing ^9^. TRPV1-positive vagal sensory neurons also innervated the lung ^29^, while CGRP-positive cells innervated the stomach ^1^ and pancreatic islets ^3^. Moreover, serotonin released from pancreatic β-cells activates 5-HT3R-expressing vagal sensory neurons ^3^, and VIP-positive neurons respond to intestinal stretch ^10^.

It has been shown that sex-dependent regulation of energy balance was significantly influenced by the quantity and activity of hypothalamic neurons involved in feeding regulation. Female mice exhibit a greater number and higher activity of anorexigenic proopiomelanocortin (POMC) neurons, which helps limit the development of obesity in females ^31^. In this regard, the uneven distribution of neurons in certain clusters between males and females could lead to sexual dimorphism in the energy balance and metabolic homeostasis. Interestingly, VN4 appeared to have a higher expression of the fibroblast growth factor receptor 1 (*Fgfr1*) gene. Fibroblast growth factor 21 (FGF21), a peptide regulated by nutrient availability ^32^, played a role in controlling hepatic gluconeogenesis and insulin sensitivity through FGF21 and FGFR1 signaling ^33^. In addition, VN4 and VN6 showed elevated levels of lysophosphatidic acid (LPA) receptor 1 (*Lpar1*). Circulating LPA levels were influenced by feeding and were higher under obesogenic conditions in animal models ^34^. The administration of LPA has been shown to have a detrimental effect on glucose homeostasis in both normal and obese mice ^35^. VN7 was found to have elevated levels of carboxypeptidase E (CPE), an enzyme responsible for processing prohormones. Hyperglycemia was observed in CPE knockout mice, with female mice exhibiting more pronounced glucose intolerance than their male counterparts ^36^. VN11, on the other hand, exhibited enhanced expression of estrogen receptor 1 (ESR1), which plays a role in regulating sex-dependent metabolic balance ^15,20,21^. While the involvement of vagal sensory neurons in these previous findings is yet unknown, it is possible that differences in gene expression patterns in vagal sensory neurons between the sexes may contribute to the sexual dimorphism in metabolic homeostasis. Hence, further studies are necessary to explore these crucial issues and to examine the physiological consequences of differences in transcriptomic profiles of vagal sensory neurons on energy balance and metabolic homeostasis.

## Research design and Methods

### Animals

All mouse care and experimental procedures were approved by the Institutional Animal Care Research Advisory Committee of Albert Einstein College of Medicine. All experiments were performed in accordance with relevant guidelines and regulations. This study is reported in accordance with ARRIVE guidelines. Eight-nine-week-old Rosa26-floxed-STOP-Cas9-eGFP mice (stock# 024857) were purchased from Jackson Laboratory. Mice were housed in cages at a controlled temperature (22 °C) with a 12:12 h light-dark cycle and fed a standard chow diet with water provided *ad libitum*.

### Single-nucleus RNA sequencing (snRNA-Seq)

Nuclei isolation and single-nucleus RNA sequencing were performed by the Singulomics Corporation (Singulomics.com, Bronx, NY, USA). Mice were euthanized with overdose of isoflurane (5% or greater). Thirty vagus nerve ganglia were collected from eight male and seven female Rosa26-eGFP^f^ mice injected with AAVrg-Cre into the liver four weeks after viral injection. Immediately after tissue collection, 16 vagus nerve ganglia from males and 14 nodose ganglia from females were flash-frozen. Tissue was homogenized and lysed with Triton X-100 in RNase-free water for nuclei isolation. The isolated nuclei were purified, centrifuged, resuspended in PBS with RNase Inhibitor, and diluted to 700 nuclei/µl for standardized 10x capture and library preparation protocol using 10x Genomics Chromium Next GEM 3’ Single Cell Reagent kits v3.1 (10x Genomics, Pleasanton, CA). Libraries were sequenced using an Illumina NovaSeq 6000 (Illumina, San Diego, CA, USA).

The libraries were sequenced with ∼200 million PE150 reads per sample on Illumina NovaSeq. The raw sequencing reads were analyzed with the mouse reference genome (mm10) with the addition of the human growth hormone poly(A) (hgGH-poly(A)) transgene sequence found in AAVrg-Cre using Cell Ranger v7.1.0. Results containing this information have been submitted elsewhere. Introns were included in the analyses. To further clean the data, Debris Identification using the Expectation Maximization (DIEM) program ^37^ was used to filter the barcodes (from raw_feature_bc_matrix of each sample), keeping barcodes with debris scores < 0.5, number of features (genes) > 250, and UMI count > 1000.

UMAP projection was generated separately and then loaded onto the Loupe Browser for downstream analysis. The UMAP projection was produced through Harmony integration, which involved using a Seurat object that incorporated normalized data with selected variable features and principal components ^38^. Initially, each sample underwent sctransform v2 normalization to identify integration features using Seurat. After merging the normalized objects, the integration features were manually selected, and 20 principal components were computed. Harmony integration aimed to unify data from different samples to improve the quality of the downstream analysis. The Harmony algorithm then facilitated the creation of a UMAP visualization, set with ‘harmony’ reduction and dimensions 1 to 20. Louvain clustering followed, applying ‘harmony’ reduction within the FindNeighbors function and the FindClusters function at a 0.5 resolution, all within the Seurat framework. Differential gene expression was determined using the 10x Genomics Loupe Browser (version 7). The significant gene test in the Loupe Cell Browser replicated the differential expression analysis in Cell Ranger.

### Statistics

To determine differential gene expression, statistical tests in Cell Ranger generated *p*-values that were adjusted for multiple testing using the Benjamini-Hochberg procedure to control the false discovery rate (FDR). To identify and visualize enriched GO terms, the web-based application Gorilla was used ^25^. Fisher’s exact test was used to determine if there was a significant association between the number of cells in the same cluster (GraphPad, Prism 10). Data were considered significantly different if the P-value was less than 0.05.

## General

I thank Drs. Woohyun Jo and Jiyeon Hwang for collecting vagus nerve ganglia.

## Author contributions

Y.H.J. performed experiments, analyzed the results, and wrote the manuscript.

## Data Availability

All data generated or analyzed during this study are included in this published article (and its Supplementary Information files).

