## Supplementary figure 1 for "Differential transcriptional profiles of vagal sensory neurons in female and male mice"

Young-Hwan Jo^1,2,3*^

**Affiliations**

^1^The Fleischer Institute for Diabetes and Metabolism

^2^Division of Endocrinology, Department of Medicine

^3^Department of Molecular Pharmacology, Albert Einstein College of Medicine, NY, USA

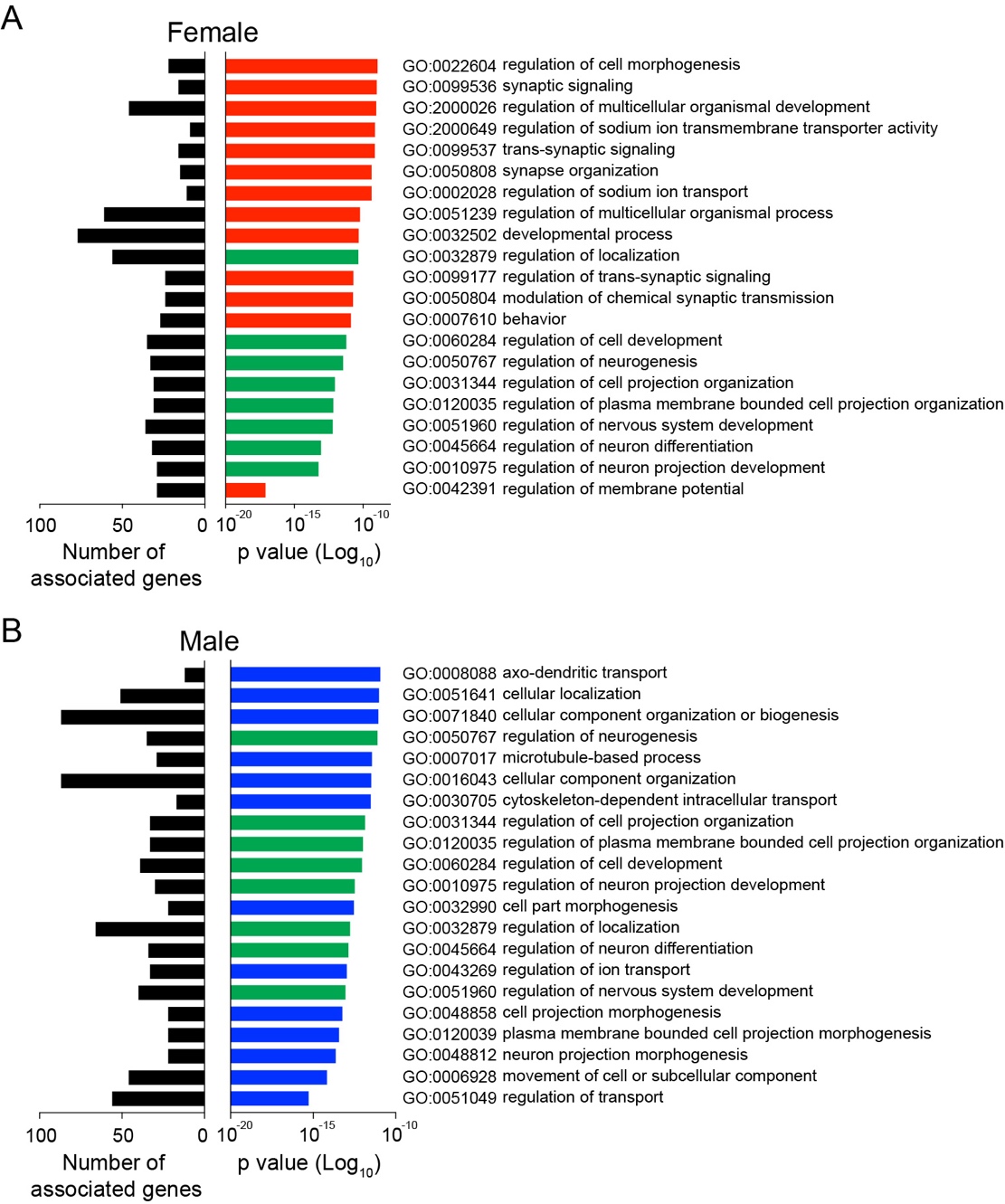

**Supplementary Figure 1**. Top 20 enriched GO terms in biological process in females and males.

**A**. Plots displaying the top 20 GO terms in biological process (left) and the number of genes associated with each GO terms in females. Green bars represent common GO terms between females and males.

**B**. Plots displaying the top 20 GO terms in biological process (left) and the number of genes associated with each GO terms in males. Green bars represent common GO terms between females and males.

**Supplementary Table 1**. Differential gene expression across the 20 clusters.

**Supplementary Table 2**. Differential gene expression among the SC1, SC2, and SCH clusters.

**Supplementary Table 3.** Differential gene expression between females and males.

**Supplementary Table 4**. Differential gene expression in the 13 neuronal clusters.
